# Stem cell intrinsic, Seven-up-triggered temporal factor gradients diversify intermediate neural progenitors

**DOI:** 10.1101/121863

**Authors:** Qingzhong Ren, Ching-Po Yang, Zhiyong Liu, Ken Sugino, Kent Mok, Yisheng He, Tzumin Lee

## Abstract

*Drosophila* type II neuroblasts produce numerous neurons and glia due to transiently amplifying, intermediate neural progenitors (INP). Consecutively born INPs produce morphologically distinct progeny, presumably due to temporal patterning in type II neuroblasts. We therefore profiled type II neuroblasts’ transcriptome across time. Our results reveal opposing temporal gradients of Imp and Syp RNA-binding proteins (descending and ascending, respectively). Maintaining Imp expression throughout brain development expands the number of neurons/glia with early temporal fate at the expense of cells with late fate. Conversely, precocious upregulation of Syp reduces the number of cells with early fate. Further, we reveal that the transcription factor, Seven-up initiates progression of the Imp/Syp gradients. Interestingly, genetic manipulations that fix Imp or Syp levels still yield progeny with a small range of early fates. We propose that the Seven-up-initiated Imp/Syp gradients create coarse temporal windows within type II neuroblasts to pattern INPs, which subsequently undergo fine-tuned subtemporal patterning.

## Introduction

Neural stem cells are heterogeneous and exhibit lineage identity as well as time-dependent developmental fate (referred to as temporal fate) [1-3]. *Drosophila* neural stem cells, called neuroblasts (NBs), acquire distinct lineage identities during early embryonic spatial patterning [4, 5]. Then, the diverse NBs likely use temporal fating mechanisms to expand neuronal diversity [1, 6-9]. Studies on *Drosophila* NBs have revealed two basic temporal fating mechanisms involved in diverse neuronal lineages at different developmental stages.

Cascades of temporal transcription factors (tTFs) underlie the specification of short series of distinct neuronal types [1]. Variants of the Hunchback-Kruppel-Nubbin and pdm2-Castor (Hb-Kr-Pdm-Cas) cascade are best known and expressed in most embryonic NBs, which make relatively short neuronal lineages before quiescence [10, 11]. Distinct tTFs are inherited by neuronal precursors born at different times, which therefore acquire different birth order-dependent cell fates[10, 11]. Similar mechanisms with variant tTF cascades guide late larval neurogenesis by different populations of medulla NBs for making the adult optic lobe (OL) [12].

One challenge in tTF cascades is producing sufficient neuronal fates for long-lived proliferative NBs. Three potential mechanisms would allow specification of more neuronal types than the number of tTFs: 1) The neuronal precursors born during tTF transitions may inherit mixtures of two tTFs and acquire combinatorial cell fates [12, 13]. 2) Multiple subtemporal fates may arise within a tTF expression window through feedforward gene regulatory networks [14]. 3) The same tTFs may confer neurons born in different competence windows with distinct cell fates [15-17]. Despite these possibilities, such short tTF cascades might still be insufficient for temporal patterning of the NBs in the post-embryonic central brain, such as the lateral antennal lobe NB that divides persistently until shortly after pupation and yields around forty-five types of projection neurons [18].

Gradients of proteins, including the IGF-II mRNA-binding protein (Imp) and Syncrip (Syp) RNA-binding proteins and the Chinmo BTB-zinc finger nuclear protein, work to control temporal fate in the long mushroom body (MB) lineages [19, 20]. The MB NBs express descending Imp and ascending Syp gradients, which endow neuronal precursors born at different developmental stages with distinct levels of Imp/Syp. Imp enhances while Syp suppresses the translation of Chinmo. Different levels of Chinmo proteins then specify three distinct mushroom body neuronal fates. The Imp/Syp gradients exist in other NBs, including the antennal lobe (AL) NBs, which yield many more neuronal types than the protracted MB lineages [18, 21]. Interestingly, the Imp/Syp temporal gradients show distinct profiles in the AL and MB NBs [20]. The progression of the rather shallow MB Imp/Syp gradients primarily occurs during late larval development coinciding with the MB temporal fate transitions. By contrast, the AL NBs express much steeper Imp/Syp gradients that progress steadily across larval development. It is not clear if and how the Imp/Syp gradients can specify all temporal fates in the rapidly changing AL lineages. In addition, Imp/Syp show mutual inhibition, as silencing Imp upregulates Syp and vice versa [20]. It is unknown how the Imp-versus-Syp dominance could be reversed and why the Imp-to-Syp switch occurs over different time courses in distinct NBs.

Canonical (type I) NBs produce post-mitotic neurons via budding off ganglion mother cells (GMCs). Each GMC typically divides into two neurons. Contrasting the production of GMCs directly from type I NBs, type II NBs first produce intermediate neural progenitors (INP) [22-24]. Each of the 16 type II NBs (8 per brain hemisphere) can generate a series of approximately 40 INPs [25]. Each INP can in turn produce around five GMCs, thus giving rise to a short sequence of neuronal siblings. A typical type II lineage therefore consists of ∼40 sublineages, each containing ∼10 neurons [25]. The complex type II pattern of neurogenesis mimics the production of neurons by mammalian neural stem cells (inner radial glial cells) through outer radial glial cells that, like INPs, can each bud off multiple intermediate precursors [26, 27].

Type II NBs also exhibit lineage identity and temporal fate [25, 28]. Labeling the progeny made by a type II NB (NB clone) reveals distinct lineage-characteristic clone morphology [25]. Six type II NB lineages (DM1-DM6) originate from the dorsomedial posterior brain surface; the remaining two lineages (DL1 & DL2) arise from the dorsolateral posterior brain surface. Most, but not all, type II NBs produce both neurons and glia [29, 30]. For instance, the DL1 lineage consistently yields glia lying between various OL neuropils [31]. Notably, the glial precursors derive from early larval-born INPs [28]. Single-cell lineage mapping further reveals that an INP can produce an invariant sequence of distinct sister neuron pairs, and that successive INPs generate a similar, but not identical, neuronal series [25]. A cascade of three tTFs (Dichaete, Grainy head, Eyeless) exists in most INP sublineages. Additional evidence supports the involvement of these tTFs in governing temporal fates within the rather short INP sublineages [28]. These observations illustrate temporal fate diversification along both axes of NB and INP self-renewal. As to the extended axis of type II NB self-renewal, it is not clear if a protracted tTF cascade exists or just gradients of proteins could guide the orderly derivation of variant INP sublineages by a single type II NB.

In order to resolve the temporal fating mechanisms in type II NBs, we specifically targeted type II NBs using complex genetic intersections [32]. We isolated pure type II NBs at different times across larval development for RNA sequencing (RNA-seq). This approach uncovered 81 significantly expressed and highly dynamic genes based on various selection criteria (see Experimental Procedures). Similar to findings from MB and AL type I NBs, Imp and Syp expression in the type II NBs show reciprocal changes over time (Imp: high to low; Syp: low to high). Notably, the opposing Imp/Syp gradients in type II NBs progress extremely quickly during early larval development. Genetic manipulations maintaining high Imp and minimal Syp expanded the number of INPs whose progeny adopt early temporal fates. Conversely, silencing Imp or overexpressing Syp elicited a selective loss of INP sublineages with early temporal fates. We further found that two embryonic tTFs, Castor (Cas) and Seven-up (Svp), known to be re-expressed in serial bursts in most early larval NBs [17], are critical for the initiation of Imp/Syp temporal gradients in type II NBs. Despite no progression of the Imp/Syp gradient, svp mutant clones carried progeny that evidently arose from INPs with multiple early temporal fates. We propose that Cas/Svp-triggered Imp/Syp gradients confer coarse temporal fates that might guide proper subtemporal patterning among INPs born within a given Imp/Syp temporal window.

## Results

### Temporally induced clones reveal type II NB temporal fates

Age-dependent fate changes in a protracted lineage can be appreciated by the order in which morphologically distinct neurons are born [18, 21]. The birth order and morphology of type II NB progeny were revealed by temporally induced twin-spot MARCM (TS-MARCM) clones among the eight type II lineages (Figure 1). TS-MARCM allows derivation of sister clones from a common progenitor via mitotic recombination [33]. It further enables concurrent labeling of the paired sister clones in distinct colors. We started by examining TS-MARCM as a type II NB buds off an INP. The INP clone consists of the neuronal and glial progenies of the INP (INP sublineage) and the NB clone contains all subsequently born INP sublineages.

**Figure 1.**
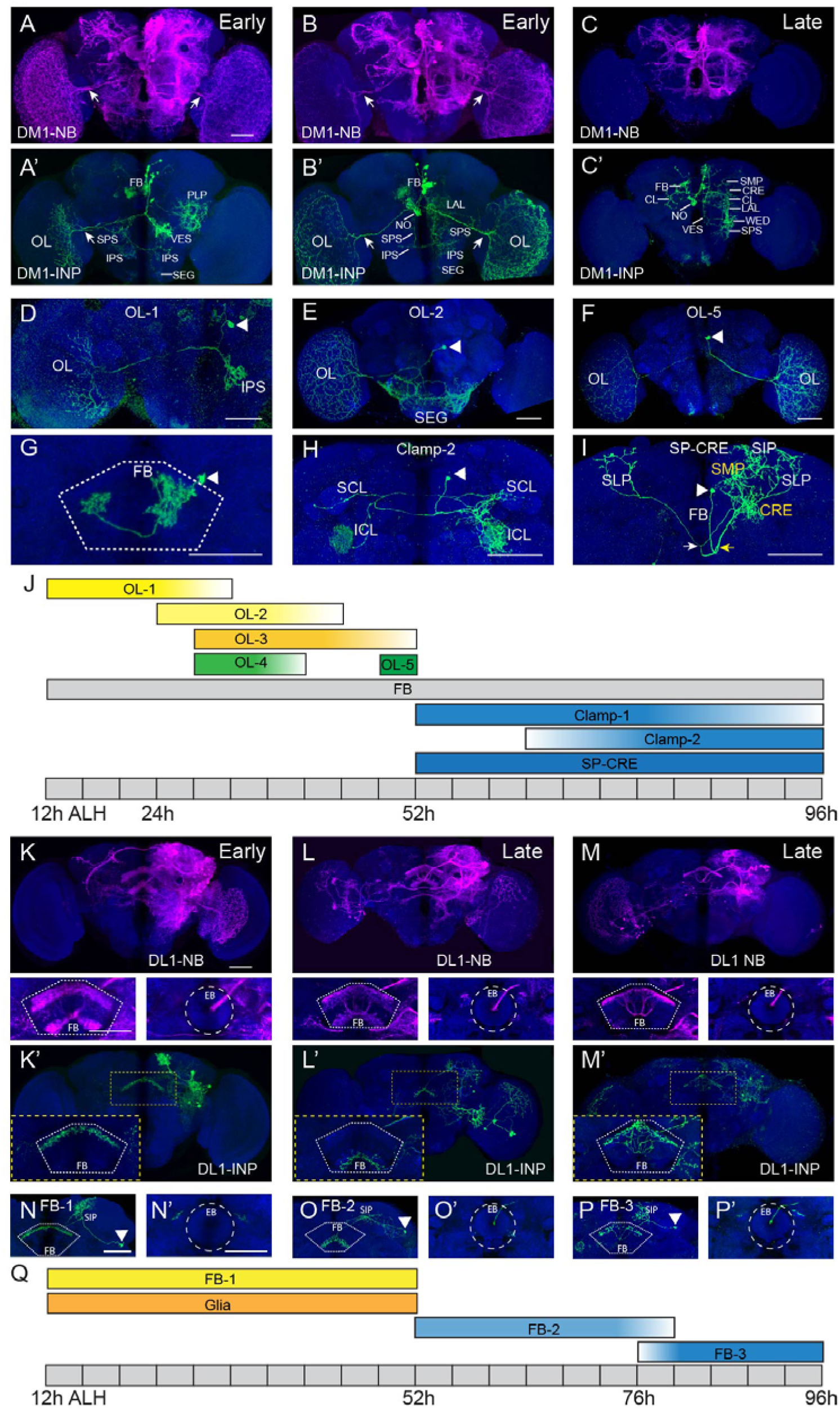
Temporal patterning in type II lineages. (A-C’) Representative confocal images of TS-MARCM NB (A-C) and INP (A’-C’) clone morphology for the DM1 lineage induced early (before 52h ALH) and late (after 52h ALH). Early NB (A, B) and INP (A’, B’) clones possess characteristic optic lobe (OL) innervations (arrows), while late NB (C) or INP (C’) clones do not. Additional brain neuropils innervated by DM1 INPs are indicated in (A’-C’). FB: fan-shaped body; SPS: superior posterior slope; IPS: inferior posterior slope; PLP: posteriorlateral protocerebrum; SEG: subesophageal ganglion; NO: noduli; VES: vest; WED: wedge; SMP: superior medial protocerebrum; CRE: crepine; CL: clamp; LAL: lateral accessory lobe. (D-I) Representative confocal images of single neurons of the OL (DF), FB (G), Clamp (H), and SP-CRE neuron classes (I). Arrowheads indicate neuronal cell bodies. Five subtypes of the OL neuron class (OL-1 to 5) innervate distinct central brain and OL areas (D-F, see Figure S1A-D for OL-3 and 4). The FB neurons (G) generally innervate two longitudinal segments of FB (white dotted region). The Clamp-2 neurons (H) form a circular commissure (named ‘the clamp commissure’) connecting bi-lateral superior and inferior clamp (SCL and ICL) regions, whereas the Clamp-1 neurons innervate bi-lateral SCL and ICL regions without forming the circular commissure (see Figure S1F). The SP-CRE neurons (I) project and bifurcate under the FB (yellow arrow). One tract continues to project anteriorly and dorsal-laterally to innervate the ipsi-lateral SMP, CRE, and LAL regions (yellow). The other tract project posteriorly and bifurcate again (white arrow). They then project anteriorly and dorsal-laterally to symmetrically innervate the bi-lateral superior protocerebrum (SP) regions (including SMP, SIP, and SLP). (J) Summary of the birth sequence of various neuronal classes of the DM1 lineage. (K-M’) Representative confocal images of TS-MARCM NB (K-M) and INP (K’-M’) clone morphology for the DL1 lineage over time. Partial projections of the FB (dotted line) and Ellipsoid body (EB, dashed circle) areas are shown in lower left and lower right panels, respectively. Insets show distinct FB innervations of INP sublineages born during early (K’, before 52h ALH) and late (L’ and M’, after 52h ALH) windows. (N-P’) Representative images of single FB neurons that first innervate superior inferior protocerebrum (SIP), and then innervate upper (FB-1, N), lower (FB-2, O), and upper (FB-3, P) FB layers, respectively. The FB-1 neurons directly project to the FB from the SIP without passing through the EB ring canal (N’). The FB-2 and FB-3 neurons project anterior-ventrally from SIP to the EB ring canal (O’-P’), and then posteriorly to innervate the FB. (Q) Summary of the birth sequence of FB neurons and glial cells of the DL1 lineage. Contaminating neurons (that overlap with neurons of interest in maximum projections) were removed from the following images (E, H, M’, P) by examining 3D images using Fluorender software. See Figure S1I, K, L, M for their original projection patterns. Scale bar: 50um.

As reported previously, an INP can produce multiple morphological classes of neurons that innervate distinct sets of neuropils [25]. Clones of a whole INP sublineage therefore show rather complex morphology (Figure 1A’-C’, K’-M’). For instance, early larval-born INP sublineages of the DM1 lineage (Figure 1A’, B’) can each yield at least three neuronal classes: (1) neurons innervating the optic lobe (OL class, Figure 1D-F and S1A-D), (2) neurons targeting the fan-shaped body (FB class, Figure 1G), and (3) neurons with elaborations in other neuropils (Posteriorlateral protocerebrum/inferior posterior slope/vest, PLP/IPS/VES in short, etc. Figure 1A’, B’). Notably, distinct sets of neuronal classes exist in late larval-induced INP clones (Figure 1C’) as evidenced by their acquisition of different trajectories and elaborations (Figure 1H-I and S1F-H). We utilized single-cell analysis of INP/GMC-derived TS-MARCM to identify late larval-born neuronal classes. In addition to the FB class (Figure 1G) that arises throughout the entire DM1 lineage, we identified two neuronal classes (Clamp class and SP-CRE class) present in most INP clones induced after 52h ALH (Figure 1C’). The Clamp-1 neurons bifurcate and innervate bi-lateral superior and inferior clamps (SCL/ICL) regions (Figure S1F), whereas Clamp-2 neurons establish a roundabout commissure (Figure S1E, the clamp commissure) that connects bilateral SCL/ICL (Figure 1H and S1G). The SP-CRE class of neurons shows elaborations in various regions of superior protocerebrum (SP) and crepine (CRE) (Figure 1I and S1H).

We could further distinguish five subtypes of DM1 OL neurons that innervate contralateral, ipsilateral, or bilateral OL regions plus various additional targets (Figure 1D-F, and S1A-D). The contralateral OL-1 subtype (Figure 1D) arises from the beginning of larval neurogenesis, while the bilateral OL-5 subtype (Figure 1F) appears last-born near 50h ALH (Figure 1J). The other three OL types (Figure 1E and S1A-D) are produced in between and show overlapping birth windows (Figure 1J). For the Clamp class of neurons, the Clamp-1 subtype emerges earlier than the Clamp-2 subtype (Figure 1J). Taken together, we demonstrate an orderly production of, not only distinct neuronal classes, but also related neuronal subtypes within a given class by the DM1 NB.

We extended similar analyses to the DL1 lineage. Examining the morphology of INP sublineages paired with NB clones reveals characteristic innervation of different FB layers over time (Figure 1K-M). Early INPs, born before 52h ALH, generate neurons that innervate the upper FB layers (Figure 1K’) directly from SP; middle INPs, born between 52 and 76h ALH, innervate lower layers (Figure 1L’) through the EB ring canal; and late INPs born after 76h ALH, innervate the upper layer with multiple branches emerging from the EB ring canal (Figure 1M’). We examined single cell morphology from INP or GMC clones and confirmed the presence of three subtypes of the FB neuron class that contribute to the FB innervation of INP sublineages (Figure 1N-P’). INP or GMC clones of these three subtypes (data not shown) also emerge sequentially during development, further supporting the sequential birth of these subtypes (Figure 1Q). In addition, we also confirmed the birth of all DL1-derived OL glial precursors before 52h ALH (data not shown; [28]).

Even in complex NB clones (Figure 1A-C, K-M), we could reliably recognize the early larval-born OL class and the late larval-born Clamp and SP-CRE classes (especially their innervation in SMP and CRE) in the DM1 lineage, as well as the early larval-born OL glia (see Figure 2G, arrow) and the late larval-born FB-2/FB-3 neurons in the DL1 lineage. Examining the NB clones of various sizes induced at different developmental times revealed a sequential disappearance of the morphological features characteristic of early-versus late-born progeny. For instance, the DM1 NB clones induced at the mid-larval stage (Figure 1C) consistently lack OL elaborations but retain the strong Clamp roundabout commissure and the dense innervations of CL/SP/CRE (Figure S1E-E’”). The absence of lower FB layer innervation (Figure 1M, lower left) in the late NB clone (Figure 1M) is consistent with the idea that INPs innervating FB lower-layer born before INPs that project to upper-layer again. Such observations further attest to type II NB temporal fate diversification and justify the use of these stage-specific neuronal/glial classes for phenotypic analysis of progeny temporal fates in genetically manipulated type II NB clones.

### Transcriptome analysis of pure type II NBs across development

We next searched for dynamically expressed genes in larval type II NBs, aiming to unravel the molecular mechanisms of temporal fate diversification in the mammalian-like neuronal lineages. We exclusively marked type II NBs via activating a conditional NB driver specifically in the NBs by genetic intersection plus subtraction [32]. RNA sequencing of type II NBs at 24h, 36h, 50h, and 84h ALH revealed 81 strong temporal dynamic genes (Figure S2A), selected based on ANOVA, fold change, and peak levels of expression (see Experimental Procedures). Many are expressed in descending or ascending gradients that extend across larval development. A few (e.g. Sip1 and br) are, however, most abundant at middle time points. We are particularly interested in those that rank top in both fold change and peak levels of expression. Based on fold change, the early-expressing E23, lin28, and Imp and the late-expressing Syp and Eip93 stand out. Among these top dynamic genes, Imp and Syp show highest peak levels of expression.

As in type I MB and AL NBs, the descending Imp and ascending Syp gradients in the larval type II NBs exhibit reciprocal changes in expression across the sampled time points (Figure S2B). However, the opposing Imp and Syp temporal gradients progress extremely quickly in type II NBs (Figure S2B). From 24h to 50h ALH, Imp decreases by 90% from ∼10,000 to ∼1000 transcripts per million (TPM) while Syp increases from minimal expression to its maximum of ∼7000 TPM. The corresponding gradients in the comparably dividing AL NBs proceed to only about 50% in the same duration. We confirmed the rapid progression in the Imp/Syp gradients of type II NBs at the protein level by immunohistochemistry, though a lag of ∼12 hours behind transcript dynamics and a continuous accumulation of Syp proteins were noted (Figure 3A-B’’’ for Imp, G-H’’’ for Syp, quantified in 3Q, R, see below).

**Figure 2.**
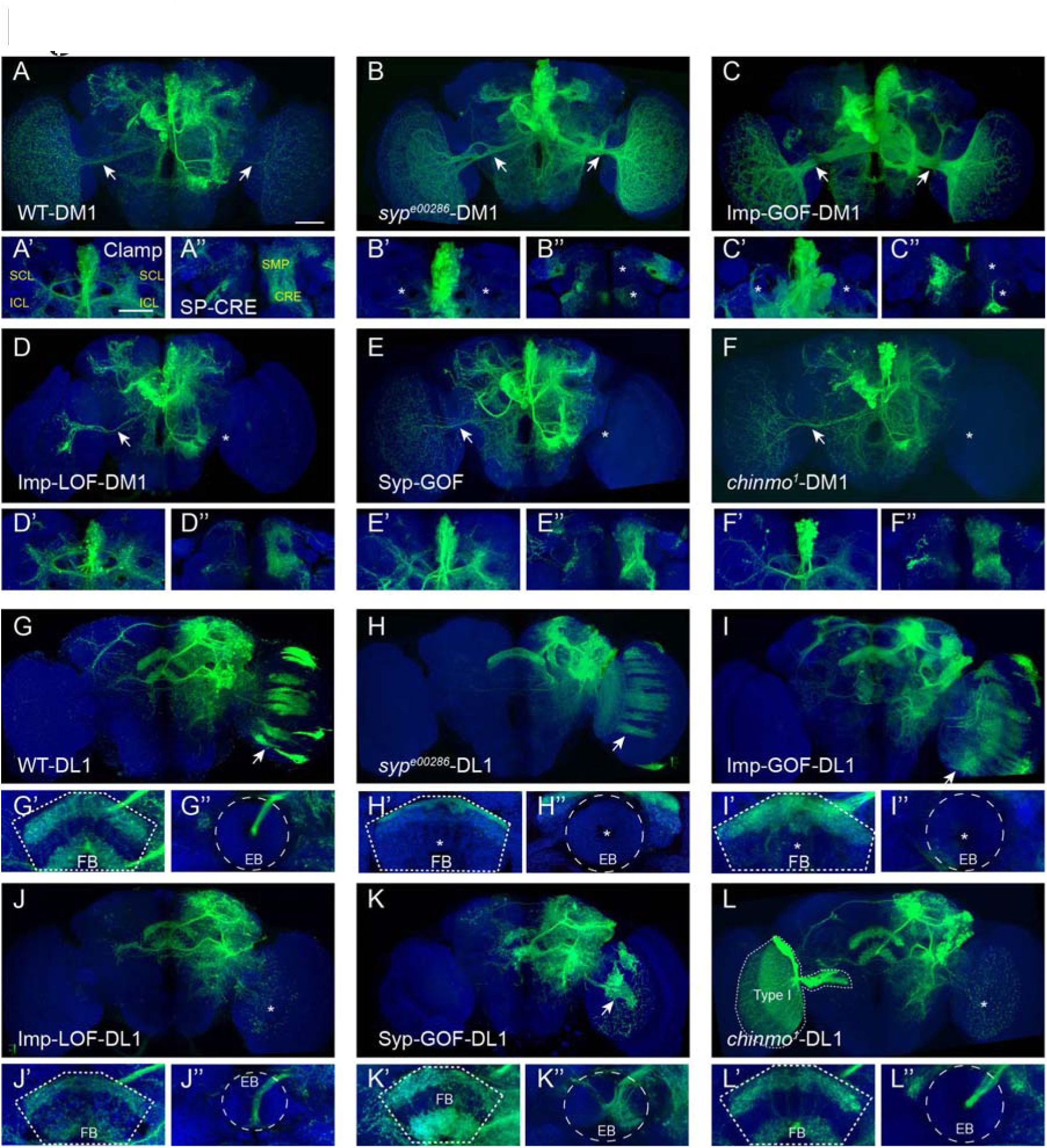
Imp, Syp and Chinmo act as temporal fate determinants. (A-F’’) Representative confocal images of MARCM clones of the DM1 lineage with altered Imp, Syp, or Chinmo expression. Compared with wild-type (WT), the OL tract (arrows) is enlarged in syp mutant (*syp^e00286^*, B) and Imp gain of function (GOF) clones (C), but reduced in Imp loss of function (LOF, via RNAi, D), Syp-GOF (E), and chinmo mutant (*chinmo^1^*, F) clones (Asterisk indicates lost). The clamp commissure (A’-F’, contributed by the Clamp-1 and 2 neurons) and SMP-CRE innervations (A’’- F’’, contributed by SP-CRE neurons) are lost in *syp^e00286^* (B’-B’’) and Imp-GOF (C’-C’’) clones (asterisks). (G-L) MARCM clones of the DL1 lineage with altered Imp, Syp, or Chinmo expression. The OL glial cells (strips indicated by arrows) are present in WT (G), *syp^e00286^*(H), and Imp-GOF clones (I, note increase), but lost in Imp-LOF(J), Syp-GOF(K, a few residual glia remain), and *chinmo^1^*(L) clones. The FB innervations and the tract to EB ring canal are shown in lower left (G’-L’) and lower right panels (G’’-L’’) respectively. The neural tract directly innervates FB upper layer in *syp^e00286^* (H’) and Imp-GOF (Γ) clones, without passing through the EB ring canal (asterisk in H’’-I’’). By contrast, the FB lower-layer innervations and the EB ring canal tract are present in Imp-LOF, Syp-GOF, and *chinmo^1^* clones. Scale bar: 50um. n≥3 samples per genotype. See also Figure S2.

**Figure 3.**
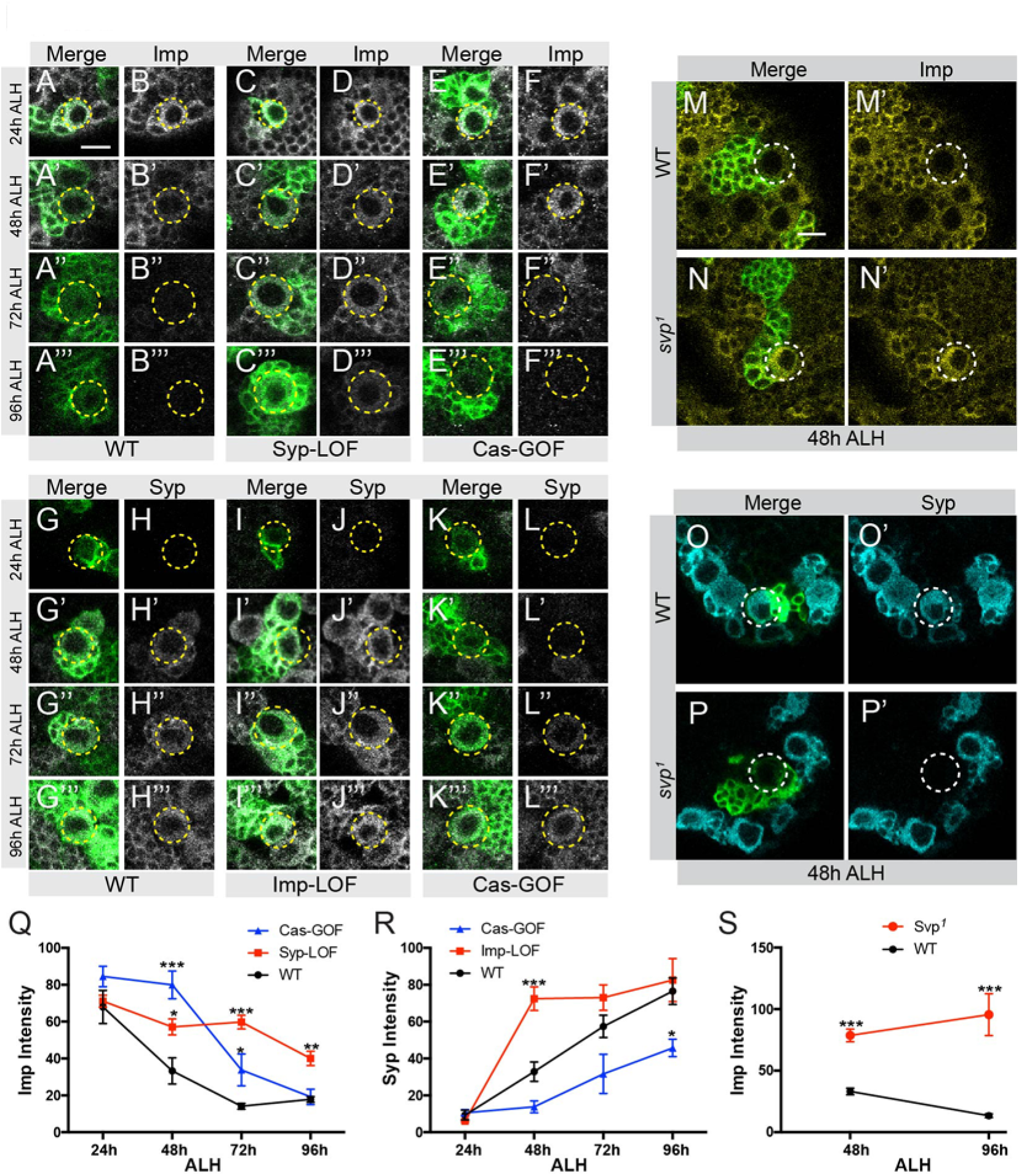
Imp/Syp repress each other and are both regulated by Cas and Svp. (A-F’’’) Imp immunostaining in DL1 NBs (yellow circles) across development (24, 48,72, and 96h ALH) in WT (A-B’’’), Syp-LOF (C-D’’’, via RNAi driven by *PntP1*-GAL4), and Cas-GOF (E-F’’’, via overexpression driven by *PntP1*-GAL4) conditions. Green: Type II lineages labeled by GFP staining. Grey: Imp staining. Imp is detectible in WT DL1 NBs only at 24 and 48h ALH. In Syp-LOF and Cas-GOF, Imp is detectible at later time points. (G-L’’’) Syp immunostainning in DL1 NBs (yellow circles) across development (24, 48, 72, and 96h ALH) in WT (G-H’’’), Imp-LOF (I-J’’’, via RNAi driven by *PntP1*-GAL4), Cas-GOF (K-L’’’) conditions. Imp-LOF accelerated Syp up-regulation, while Cas-GOF slowed down Syp up-regulation. (M-P’) Imp and Syp immunostaining in WT and *svp^1^* MARCM clones at 48h ALH (induced at 8-12h ALH). Svp loss-of-function enhanced Imp expression (M-N’) and prevented Syp emergence (O-P’). (Q-S) Quantification of Imp or Syp immunostaining (greyscale intensity) from (A-P’) (See Experimental Procedures). Data are shown as means± SEM. n≥3 samples per genotype. * indicates significant difference from WT Imp or Syp levels by one-way ANOVA (Q-R), unpaired t-test (S, 48h ALH), or Mann-Whitney test (S, 96h ALH). *: p<0.05, **: p<0.01, ***: p<0.001. Scale bar: 10um.

### Imp, Syp, and Chinmo regulate type II NB temporal fates

We probed the roles of Imp/Syp gradients in the derivation of complex type II neuronal lineages. To accomplish this, we repressed or expanded the expression of Imp or Syp in stochastic type II NB clones by MARCM or in all type II lineages via targeted RNAi followed by MARCM-based phenotypic analysis. We determined the NB temporal fate history in adult DM1 and DL1 NB clones where, despite complex clone morphology, various neuronal classes reflecting different temporal fates are recognizable (Figure 1).

Silencing Syp or enhancing Imp resulted in enlarged NB clones that were packed with progeny normally made by early-born INPs (Figure 2A-C for DM1 lineage, 2G-I for DL1 lineage). In the DM1 lineage, we observed great expansion of the early temporal fate-dependent OL-elaboration neurons (Figure 2B and C) but loss of majority of the late temporal fate-dependent Clamp commissure neurons and SP-CRE neurons (Figure 2B’-B’’, C’-C’’). In the DL1 lineage, the later two types of FB neurons are missing from the extraordinarily large NB clones (Figure 2H and I) where the numerous FB neurons exclusively target the upper FB part without passing through the EB ring canal (Figure 2H’-H’’, I’-I’’). These DM1/DL1 phenotypes indicate an overproduction of INP lineages with early temporal fates at the expense of those with late temporal fates. Our data imply that the type II NBs with persistently high Imp or minimal Syp expression have continuously yielded INPs with early temporal fates.

The opposite phenotypes were obtained following repression of Imp or misexpression of Syp (Figure 2D, E for DM1 lineage, 2J, K for DL1 lineage). The manipulated DM1 NB clones have relatively normal neuronal number (Figure 2D, E) but resemble a much later-induced NB clone (Figure 1E) with some residual OL-elaboration neurons. Analogously induced DL1 NB clones rarely contain any early-born OL glia (Figure 2J and K). By contrast, the EB canal tract and late innervations of the FB were still present in those manipulated DL1 clones (Figure 2D’-D’’, E’-E’’, J’- J’’, K’-K’’), suggesting the presence of the late-born FB-2 and FB-3 neurons. These phenomena are consistent with a selective loss of progeny with early temporal fates. Taken together, our data suggest that Imp promotes and Syp antagonizes early temporal fates and that the opposite applies to late temporal fates.

Nonetheless, perturbing type II NB temporal fate progression in the same direction by manipulating Imp versus Syp did not create identical phenotypes. For instance, Imp gain-of-function (Figure 2C and I), but not Syp loss-of-function (Figure 2B and H), caused thickened primary bundles with reduced terminal elaborations in both DM1 and DL1 NB clones, which might result from ectopic Imp activity in postmitotic cells. In addition, the DL1 dorsal midline-crossing neurons were retained in Imp or Syp overexpression (Figure 2I and K) but not detectable in Syp or Imp repression (Figure 2H and J). The dorsal midline-crossing neurons of the DL1 lineage normally arise around the mid-larval stage. Their selective retention in Imp or Syp overexpression, but not after Syp or Imp repression, might reflect the promotion of middle temporal fates by co-existence of Imp and Syp.

Moreover, the NB clones lacking late- or early-derived trajectories still show rather complex morphology (Figure 2B-E, H-K) resulting from not only diverse neurons of common INP origins but also distinct INP lineages. For instance, there exist serially derived types of OL-elaboration neurons in the Syp mutant DM1 NB clones (Figure 2B). These phenomena suggest that type II NBs without detectable Syp or Imp can still produce INPs with various early or late temporal fates.

We next examined if Chinmo [19], a known Imp/Syp downstream effector [20], governs neuronal temporal fates in type II NB lineages. Our RNA-seq recovered *chinmo* as a temporal dynamic gene with a shallow descending gradient across entire larval development (Figure S2). By immunostaining, Chinmo proteins were present in early larval type II NBs but became undetectable by 72h ALH (data not shown). Moreover, *chinmo* mutant DM1 (Figure 2F) and DL1 NB clones (Figure 2L) show a selective loss of early-derived progeny including the DM1 OL-elaboration neurons and the DL1 OL glia. Consistent with the notion that Imp promotes while Syp suppresses Chinmo translation, the Syp mutant type II NBs remained positive for Chinmo proteins throughout 96h ALH (data not shown). Taken together, our data show that the opposing Imp/Syp gradients govern type II NB temporal fates at least partly through regulating Chinmo.

### Cas/Svp govern type II NB temporal fates via controlling Imp/Syp

Imp and Syp show negative cross-regulation in the MB NBs, as repressing one enhances the other [20]. We examined if this is also true in type II NBs by targeted Imp/Syp RNAi using *PntP1*-GAL4, which drives expression in type II NBs and newly generated INPs [34]. Knockdown of Syp by RNAi led to persistent expression of Imp in larval type II NBs and their newborn progeny (Figure 3A-D’’’, quantified in 3Q). Conversely, repressing Imp greatly accelerated Syp up-regulation (Figure 3G-J’’’, quantified in 3R). This confirms cross-inhibition between Imp and Syp in type II NBs. However, loss of Syp did not completely block the drop in Imp expression from 24h to 96h ALH (Figure 3C-D’’’; quantified in Figure 3Q). Furthermore, absence of Imp did not de-repress Syp at 24h ALH (Figure 3I and J). It appears that some Imp/Syp-independent mechanism(s) govern the beginning Imp/Syp levels and their initial changes in type II NBs. The acute initial regulation followed by continual cross-inhibition may drive the prompt reversal of Imp-versus-Syp dominance in early larval type II NBs.

We hypothesized that the classic tTF cascade might regulate Imp/Syp gradient progression. RNAseq data indicate that tTFs, Cas and Svp are expressed dynamically in type II NBs (Figure S2), whereas Hb/Kr/Pdm are minimally expressed (data not shown). Cas, a zinc finger transcriptional factor, has been shown to specify late temporal fates or close temporal identity windows [13, 35-37]. Svp, a member of the nuclear hormone receptor family [38], functions as a switching factor for the Hb to Kr transition by suppressing Hb [39-41]. Beside the role of switching factor, Svp could also act as a sub-temporal factor to sub-divide a wider temporal window [42]. Bursts of Cas/Svp expression in most, but probably not all, early larval NBs have been shown to govern the progression and termination of post-embryonic neurogenesis [17, 43]. In a parallel study (Yang, et al. In preparation), we demonstrate that the opposing Imp/Syp gradients co-regulate neuronal temporal fate and NB senescence. We therefore posited whether the Cas/Svp pulse triggers the Imp/Syp gradients in type II NBs. We examined this hypothesis by first determining the requirement of Cas/Svp for proper development of the DM1 and DL1 lineages.

Knocking out svp or expressing transgenic cas in type II NBs elicited great expansion of progeny with early temporal fate at the expense of those with late temporal fate. The svp mutant MARCM DM1 NB clones carry supernumerary neurons with early temporal fates, as evidenced by exuberant OL/SEG elaborations (Figure 4A-B) and loss of clamp/SP/CRE innervations (Figure 4B’-B’’). The DL1 svp^−/−^ NB clones show densely packed OL glia (Figure S3E) but lack later types of FB neurons (Figure S3E’-E’’) despite their production of neurons in excess. Notably, the DM1 svp^−/−^ NB clones apparently contain multiple early INPs with distinct temporal fates, including INPs that innervate SEG (INPs making OL-2 neurons), INPs that innervate ipsi-lateral upper lobula (INPs making OL-4 neurons), and INPs that innervate bi-lateral OL (INPs making OL-5 neurons), etc. Expressing transgenic cas in type II NBs with *PntP1*-GAL4 (Figure 4C and Figure S3F) resulted in similar phenotype as svp mutants, except that some late clamp/SP/CRE processes were still present for DM1 clones, and that 80% of DL1 clones possess neural tracts through the EB ring canal (Figure 4C’-C’’, Figure S3F’-F’’, and data not shown). By contrast, no obvious defects could be found in the NB clones with the opposite manipulations of overexpressing svp or knocking out cas prior to larval hatching (Figure 3B-C’’, G-H’’). These results confirm the involvement of Svp/Cas in patterning type II NB temporal fates. It further implies that Svp promotes and Cas suppresses temporal fate progression.

**Figure 4.**
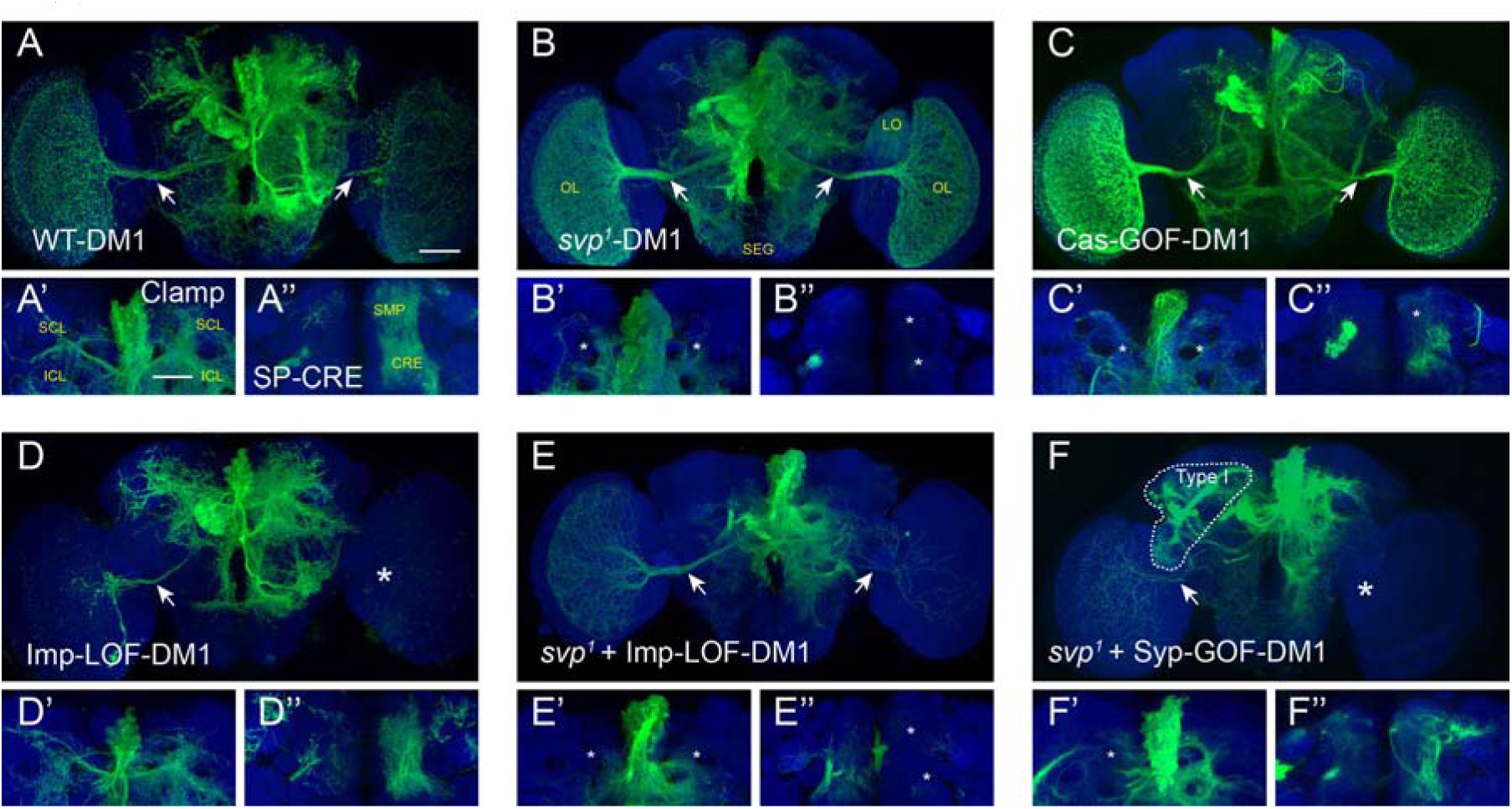
Svp and Cas regulate temporal fate specification, potentially through Imp/Syp. Adult MARCM clones of the DM1 lineage (induced at 8-12h ALH) are shown in (A-F), with the clamp commissure and SMP-CRE innervations shown in (A’-F’) and (A’’-F’’), respectively. (A-D’’) The OL tracts (arrows) are enlarged in *svp^1^* (B) and Cas-GOF (C) clones, and reduced in Imp-LOF clones (D). The clamp commissure and SMP-CRE innervations are absent (asterisks) in *svp^1^* clones (B’-B’’), reduced in Cas-GOF clones (C’-C’’). The *svp^1^* clone (B) includes innervations of bi-lateral OL, SEG, and ispi-lateral upper lobula (LO) regions. (E-F’’) Reducing Imp or enhancing Syp expression in *svp^1^* clones reduced the OL tracts (arrows in E-F, asterisk in F). Reducing Imp expression in *svp^1^* clones did not restore the clamp commissure or SP-CRE innervations (E’-E’’). Enhancing Syp expression in *svp^1^* clones restored some innervations of the SMP-CRE (F’-F’’). The dotted area in (F) is a contaminating type I lineage clone. n≥3 samples per genotype. Scale bar: 50um. See also Figure S3.

We next examined Imp and Syp levels in svp mutant NB clones at 48h and 96h ALH. Within the same mosaic brains, wild-type NBs had minimal Imp and strong Syp expression (Figure 3M-M’, O-O’), whereas homozygous svp mutant NBs had high Imp and undetectable Syp expression (Figure 3N-N’, P-P’). We quantified the Imp immunostaining and detected no progression of Imp/Syp gradients in svp mutant type II NBs (Figure 3S). These data are consistent with a recent study showing persistent Imp expression in svp mutant NBs [43]. We also examined Imp/Syp levels in type II NBs with continuous Cas expression. Notably, Cas overexpression delayed the onset of Imp/Syp gradient progression, but failed to maintain high-Imp/low-Syp levels indefinitely (Figure 3E-F’’’, K-L’’’, quantified in Figure 3Q-R). We detected minimal progression by 48h ALH, but substantial yet variable progression at 72h ALH. The delayed onset of the Imp-to-Syp transition may explain why the Cas-overexpressing DM1 NB clones contain some progeny with late temporal fate in addition to supernumerary cells with early temporal fate (Figure 4C-C’’). Moreover, about 80% of Cas-overexpressing DL1 NB clones have acquired a weak neurite track passing through the EB ring canal (data not shown), indicating presence of a small number of late-born FB-2 or FB-3 neurons.

If the svp mutant phenotype is caused by NBs with persistent high levels of Imp and no Syp, one should be able to suppress or even reverse the phenotype by repressing Imp or overexpressing Syp. Indeed, silencing Imp (Figure 4E, DM1; Figure S3J, DL1) or continuously expressing Syp (Figure 4F, DM1; Figure S3L, DL1) reversed the svp mutant phenotype to a certain extent. Reducing Imp expression reduced early OL projections (Figure 4E), but did not restore the late clamp and SP-CRE projections in the DM1 lineage (Figure 4E’, E’’). Over-expressing Syp in Svp mutant also reduced early OL projections (Figure 4F) and restored some innervations in the SP-CRE region (Figure 4F’’), although no apparent clamp innervation was observed (Figure 4F’). These two manipulations also reduced early glial cells (Figure S3J-L) and restored the EB canal tract and projections from the EB canal to the FB upper layer (Figure S3J’-J’’, L’-L’’). Taken together, the Cas/Svp bursts in post-embryonic NBs somehow jumpstart the Imp/Syp gradients that cell autonomously govern type II NB temporal fates.

### A sequence of Cas>Svp expression precedes Imp-to-Syp transition

It has been shown in larval thoracic NBs that the timing of Cas and Svp bursts varies from one to another NB but Cas expression mostly precedes Svp expression [17]. To examine if type II NBs exhibit similar temporal dynamics, we mapped the expression of Cas/Svp in early larval type II NBs by immunostaining. Cas could be detected in various subsets of type II NBs in different brains from 6-30h ALH (Figure 5A). Compared to Cas, Svp appeared in smaller various subsets and at slightly later time points. To control for lineage variations, we examined the dynamics of Cas and Svp expression specifically in the DM1 and DL1 NBs. We clearly detected Cas prior to Svp in both lineages and noticed a slightly earlier appearance of the Cas/Svp pulse in the DL1 NB (Figure 5B-C).

**Figure 5.**
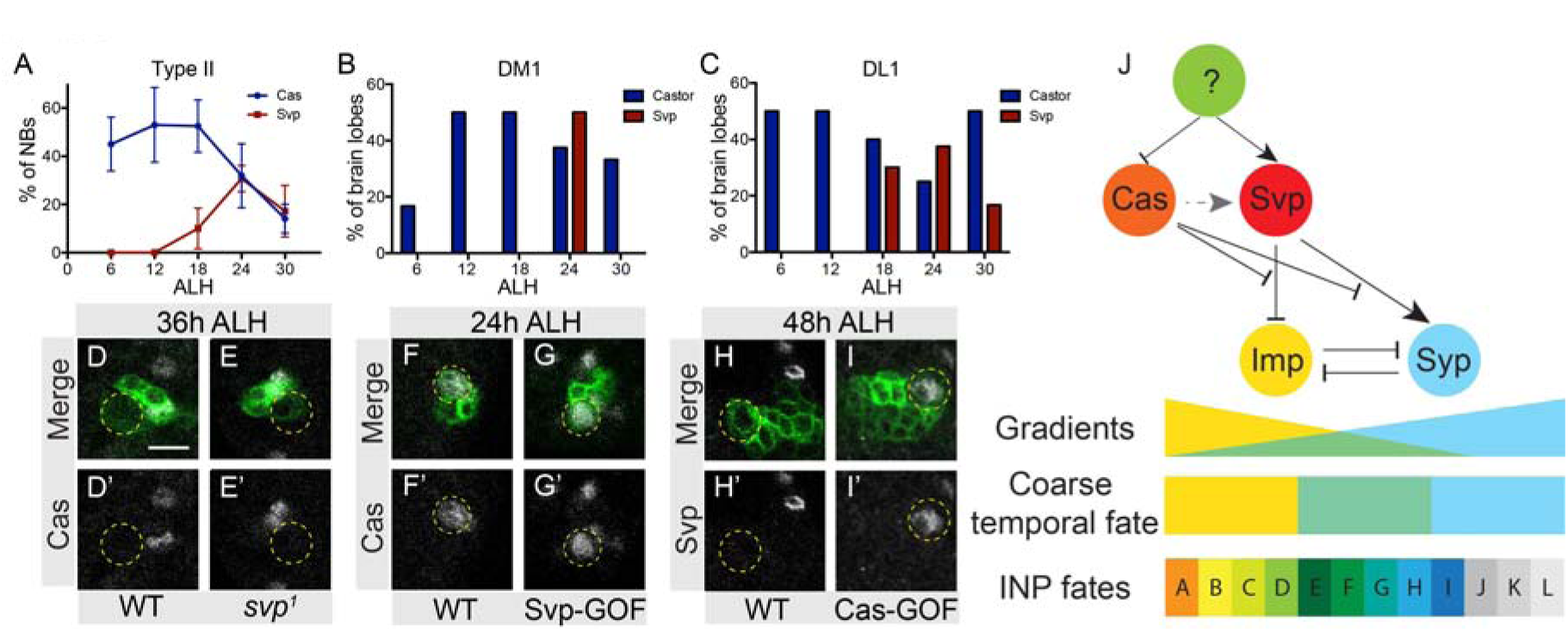
Cas>Svp expression precedes Imp-to-Syp transition. (A) Mean percentage of Cas+ and Svp+ NBs among the eight type II NBs. Cas appears first while Svp arises later, and both disappear after 30h ALH. (B and C) Percentage of brain lobes possessing Cas+ and Svp+ DM1 and DL1 NBs. (D-E) Loss of Svp did not extend the Cas expression into later time windows. (F-G) Ectopic expression of Svp did not suppress Cas expression at 24h ALH. (H-I) Ectopic expression of Cas extended Svp expression into latter time window (48h ALH). (J) A working model of temporal fate specification in type II NBs. A brief burst of Svp suppresses Imp and activates Syp, thus enabling the progression of Imp/Syp gradients. The relative level of Imp versus Syp specifies coarse temporal windows. Then, subtemporal pattering mechanisms subdivide each coarse window into smaller temporal windows to specify distinct INPs fates. Cas can promote, but is not necessary for, Svp. Thus, the repression of Cas and emergence of Svp seem to be under the control of an unknown temporal cue (question mark). n≥3 samples per genotype. Scale bar: 10um.

Given the temporal coupling of Cas and Svp bursts, we wondered if Cas and Svp regulate each other’s dynamic expression. Knocking out svp did not affect the post-embryonic burst of Cas (Figure 5D-E’); neither did induction of transgenic svp (Figure 5F-G’). However, the transient expression of Svp could be drastically prolonged in type II NBs that continuously express Cas (Figure 5H-I’). This indicates that ectopic Cas extend Svp expression but paradoxically antagonizes the activation of Imp/Syp gradients by Svp. Nonetheless, the endogenous Cas bursts appear dispensable for endogenous Svp expression in type II NBs, as the embryonic-induced cas mutant NB clones (Figure S3C-C’’, H-H’’) are grossly normal, sharply contrasting the greatly enlarged svp mutant clones that carry only progeny of early temporal fates (Figure 4B and S3E). Taken together, we propose that the timely repression of Cas and transient activation of Svp occur independently to jumpstart Imp/Syp temporal gradients (Figure 5J).

## Discussion

### Imp/Syp gradients govern type II NB temporal fates

Our RNA-seq of pure *Drosophila* type II NBs across larval development revealed 81 strong temporal dynamic genes, which include previously reported temporal factors Imp and Syp [20]. The opposing Imp/Syp temporal gradients are shallow and progress minimally until late larval development in the MB NBs. The same temporal gradients in the AL NBs are steep and progress steadily throughout pre-wandering larval development. Notably, type II NBs show further steep Imp/Syp mRNA gradients with complete reversal of Imp-versus-Syp dominance by the beginning of 3^rd^ instar. We identified various neuronal classes characteristic of early or late larval-born INP sublineages within the DM1 and DL1 lineages. We could greatly increase the number of progeny that the early INPs produce by enhancing Imp or suppressing Syp, or selectively deplete early progeny by repressing Imp or overexpressing Syp. In general, the opposing Imp/Syp gradients govern type II lineage temporal fate determination. However, we are not sure if absolute levels of Imp and/or Syp or their relative levels confer specific temporal fates, and how a given Imp/Syp level is perceived by individual GMCs of the same INP origin. More sophisticated controls over Imp and Syp levels and finer temporal fate readouts are required to resolve these details. Nonetheless, type II NBs locked at the initial Imp/Syp levels still underwent some limited temporal fate progression, as best evidenced by the presence of various OL-elaboration neurons in the svp mutant DM1 NB clones (Figure 4B). We therefore propose that the rapid Imp/Syp gradients confer type II NBs with coarse temporal fates, which guide or permit subtemporal patterning among INPs born within a given Imp/Syp temporal window (Figure 5J).

Chinmo, a known Imp/Syp downstream effector, promotes early temporal fates in diverse neuronal lineages. Consistent with Imp promoting and Syp suppressing Chinmo translation, Chinmo proteins could be detected in type II NBs only when Imp dominates Syp. Maintaining high Imp and minimal Syp extended the expression of Chinmo proteins into late-larval type II NBs. As expected, chinmo mutant type II NB clones show a selective loss of early INPs’ progenies, including the OL-elaboration neurons in the DM1 lineage and the OL glia in the DL1 lineage. The same phenotypes exist in Imp-depleted or Syp-enhanced type II NB clones. These phenomena implicate Chinmo as a main Imp/Syp downstream temporal factor in the complex type II NB lineages. However, chinmo mutant NB clones appear more exuberant than Imp/Syp-manipulated clones, as evidenced by their possession of more widespread terminal elaborations. It is possible that Imp/Syp post-transcriptionally regulate additional temporal genes, critical for Chinmo-independent temporal fate specification and/or neuronal differentiation.

Temporal fate diversification continues in type II NBs during late larval development with full Syp expression and no detectable Imp. Extensive subtemporal patterning mechanisms are needed for refinement of temporal fates among later-born INPs. Extrinsic inputs could help govern the subtemporal patterning of late larval neurogenesis and simultaneously coordinate brain development with overall organism growth. Interestingly, the ecdysone-induced protein Eip93 was recovered as a late temporal gene in diverse NBs, including type II NBs ([20]; Figure S2). Moreover, Doe lab demonstrates in a parallel study (Syed MH, Mark B, and Doe CQ, personal communication) that the dynamic expressions of Eip93 and other late temporal factors depend on ecdysone signaling. Blocking the extrinsic input compromised the expression of various late temporal genes. Taken together, the intrinsic Imp/Syp temporal gradients act in combination with extrinsic temporal factors to diversify type II NB temporal fates across protracted larval development.

### Cas/Svp bursts trigger the Imp-to-Syp switch

A brief series of Cas/Svp bursts initiates the prompt switch of Imp-versus-Syp dominance in type II NBs. Cas likely precedes Svp in their post-embryonic reexpression. But the dynamic expressions of Cas and Svp appear independently yet simultaneously controlled by an unknown temporal cue (Figure 5J). Furthermore, Cas is dispensable and ectopic Cas delays the onset of Imp/Syp gradients. It seems that Cas represses while Svp promotes the switch of Imp-versus-Syp dominance. Precocious and continuous Svp expression did not accelerate or alter the Imp/Syp gradients, further implicating involvement of unknown temporal cues not only in the regulation of dynamic Cas/Svp expressions but also in confining the acute Svp action (Figure 5J).

It is unclear how a transient Svp activity can drive the Imp-to-Syp transition over a course of ∼24 hours. Observing how the Imp or Syp levels change in the absence of the other (thus eliminating their mutual inhibition) should provide some clues.

Notably, Imp still dropped significantly from 24-96h ALH in the Syp-depleted NBs. We can, therefore ascribe the initial drop in Imp to Svp and the final full repression of Imp by Syp. As to Syp, loss of Imp failed to de-repress Syp at 24h ALH but accelerated the upregulation of Syp by some Imp-independent activity. It seems that the absence of Syp right after NB quiescence is not due to high Imp, and that Syp is possibly upregulated in early larvae by Svp. We therefore speculate that the acute Svp signal can repress Imp as well as promote Syp and ultimately place Syp over Imp in their winner-take-all competition (Figure 5J). Both timing and intensity of the Cas/Svp bursts may vary among different NBs, which can potentially shape distinct Imp/Syp gradients in different neuronal lineages. To address this requires monitoring and manipulating Cas/Svp expressions as well as their dynamic activities in identifiable NBs that show distinct profiles of Imp/Syp gradients.

### Combinatorial temporal fate specification in INP progenies

The Imp/Syp gradients in type II NBs endow INPs born at different times with distinct levels of Imp/Syp proteins, which is required to confer distinct INP identity. Each INP further expresses a cascade of tTFs to assign its serially derived GMCs with distinct cell fates [28]. Imp/Syp and their downstream effectors could interact with the INP tTFs to specify terminal temporal fates in post-mitotic cells. Multiple feedforward gene regulatory loops might also be involved to control the expression of terminal selector genes [14]. Distinct Imp/Syp levels may specify different neuronal classes among the GMCs of the same birth order that inherit the same tTF from INPs. It is not possible to resolve such complex temporal fating mechanisms without further sophisticated single-cell lineage mapping tools.

### Conserved temporal factors in mammalian neural stem cells

Converging evidence indicate that conserved tTFs control neuronal diversity from *Drosophila* to mammalian species [7-9, 28, 44]. Mammalian neural stem cells show temporally patterned neurogenesis as in the developing fly brain. The Ikaros family zinc finger 1, orthologues of *Drosophila* Hb, specify early temporal fate in both the retina and the cortex [45, 46]. A recent study showed that an ortholog of *Drosophila* castor, Casz1, promotes late neuronal fates in the mouse retina [47]. The Chick Ovalbumin Upstream Promoter-Transcription Factors (COUP-TFI and II), mammalian orthologues of *Drosophila* Svp, promote the temporal transition from neurogenesis to gliogenesis in neural stem/progenitor cells [48]. COUP-TFI also orchestrates the serial generation of distinct types of cortical interneurons [49]. Loss of COUP-TFs resulted in over-production of early-born neuronal fates, at the expense of late-born glial and interneuron fates. Thus, the COUP-TFs seems to be functionally conserved to *Drosophila* Svp.

Notably, a descending Imp gradient exists in mouse neural stem cells and governs temporal changes in stem cell properties [50]. Given the use of INPs in both type II NB lineages and mammalian neurogenesis, it would be interesting to determine if homologs of Imp/Syp/Chinmo play analogous roles in regulating the temporal fates of mammalian neural stem cells, and if these genes mediate the effects of the conserved tTFs.

## Experimental Procedures

### Fly stocks

Flies were reared at 25°C on standard medium. For stocks made in this study, the complete nucleotide sequences of the plasmids will be provided upon request. See Supplemental Experimental Procedures for the complete list of *Drosophila* stocks utilized and genetic crossing schemes for each figure.

### Clonal analysis

Newly hatched larvae were collected over a 4h window, and put into vials (100 larvae/vial) containing fly food. For temporal mapping of distinct INP sublineages (Figure 1), larvae were heat-shocked (37°C) at successive 4h windows from 12h to 88h ALH. Over the course of larval development, the NB clone frequency reduced over time. Thus, a second round of mapping for late larval development (from 58h to 96h ALH) was performed. The heat-shock durations were titrated specific for each time window based on experience. To induce adult MARCM clones with manipulated Imp, Syp, Chinmo, or Svp expression, larvae were heat-shocked 14min at 8-12h ALH. For analysis of larval mutant MARCM clones, the heat-shock time were increased to 1 hour. For induction of *cas^24^* adult MARCM clone, progeny were heat-shocked 14min at 4-8h after egg laying. All flies were returned to 25°C after heat-shock.

### Immunohistochemistry and Confocal microscopy

Brain samples of larvae or 3-6d old adult were dissected in ice-cold phosphate-buffered saline (PBS) and fixed in 2% paraformaldehyde (PFA) for 50min at room temperature. After two rinses in PBT (PBS plus 0.5% Triton X-100), they were blocked for 1 hour in blocking solution (PBT containing 4% Normal Goat Serum). Then, samples were incubated overnight in blocking solution containing primary antibodies at 4 degree. After four 20-min wash in PBT, secondary antibodies were added and again incubated overnight. Brains were transferred into PBS and mounted in slowfade (ThermoFisher Scientific) after four 20-min wash in PBT to remove secondary antibodies. The primary antibodies include: Chicken anti-GFP (1:1000; Life Technology, A10262), Rabbit anti-GFP (1:1000; Life Technology, A11122), Rat anti-mCD8 (1:1; Life Technology, MCD0800), Rabbit anti-DsRed (1:500; Clonetech, #632496), Rat anti-Deadpan (1: 100; Abcam, ab195173), Mouse anti-nc82 (1:50; Developmental Studies Hydridoma Bank or DSHB), Rabbit anti-Chinmo (1:1000;[19]), Rabbit anti-Imp (1:600; Paul Macdonald, University of Texas at Austin;[51]), Guinea pig anti-Syp (1:500; Ilan Davis, Univeristy of Oxford; [52]), Mouse anti-Svp (1:10, DSHB 5B11;[39]), Rabbit anti-Castor (1:500; Ward Odenwald, NIH;[36]), Mouse anti-Broadcore (1:100; Gregory Guild, University of Pennsylvania;[53]). Corresponding secondary antibodies were from Jackson ImmunoResearch (West Grove, PA). Image stacks of whole-mount fly brains were taken using a Zeiss LSM 710 confocal microscope and processed with Adobe PhotoShop. To compare Imp/Syp levels across various genotypes, images were taken using the same confocal setting (pinhole size, gain, laser power, etc).

### RNA-seq and Data analysis

The methods for genetic labeling and sorting of type II NBs, as well as subsequent RNA extraction, Library preparation, sequencing, and dynamic gene analysis were described previously[20, 32]. Here we extend RNA-Seq of type II NBs to multiple time points during development (24h, 36h, 50h, and 84h ALH), to identify dynamically changing genes. Eighty-one temporal dynamic genes were recovered as they meet either of the two sets of selection criteria. The first set of criteria: 1) the p-value (one-way ANOVA) for comparison between TPM values of the four time points is less than 0.05. 2) the peak TPM value is larger than 120. 3) the maximum fold change of the average TPMs in pair-wise comparisons is bigger than 6. The second set of criteria: 1) the p-value (one-way ANOVA) is less than 0.001. 2) the peak TPM value is larger than 120. 3) the maximum fold change is bigger than 3. The RNA-Seq data will be deposited in NCBI Gene Expression Omnibus.

### Statistical Analysis

To quantify NB Imp/Syp protein levels, the gray-scale intensities (from 0-255) of areas of interest (circled regions in Figure 3) were quantified using Image J software (National Institute of Health). To compare Imp/Syp levels between WT, Imp/Syp-LOF, Cas-GOF conditions, we employed one-way ANOVA followed by planned pair-wise comparisons with Dunnett’s *post hoc* test. For comparison of Imp levels between WT and *svp^1^* clones, we employed unpaired t-test or Mann-Whitney test. All tests in this study were two-sided. Statistical tests were done using Prism 6 (Graphpad Software). Asterisks indicate levels of significant differences (*: p<0.05, **: p<0.01, ***: p<0.001).

## Author Contributions

Q.R. and T.L. designed the study. Q.R., K.M., and Y.H. performed experiment. C.Y., Z.L., and S.K. performed RNA-Seq experiments. Q.R. and S.K. analyzed the data. Q.R. and T.L. wrote the manuscript.

## Acknowledgements

We thank Janelia Workstation project team and Flycore for technical assistance. We are grateful to Paul Macdonald, Ilan Davis, Ward Odenwald, and Gregory Guild for valuable antibodies. We thank Stefan Thor, Bloomington Drosophila Stock Center, Vienna Drosophila RNAi center, Transgenic RNAi project at Harvard Medical School, and Drosophila Studies Hydridoma Bank for essential stocks and reagents. We thank Chris Doe for sharing unpublished results on regulation of temporal patterning by ecdysone signaling and critical reading of the manuscript, Rosa Miyares for inputs, and Crystal Di Pietro for administrative support. This work was funded by Howard Hughes Medical Institute.

